# AOPTK: Open-Source Software for AI-Assisted Toxicological Literature Analysis

**DOI:** 10.64898/2026.10.29.761316

**Authors:** Robin Durník, Lola Bajard, Luděk Bláha, Pavel Babica, Iva Sovadinová, Helge Hecht

## Abstract

**Motivation:** Toxicologists today face a challenge: although vast amounts of data describing how chemicals affect human health are available across scientific publications and regulatory reports, much of this information remains underutilized because extracting and interpreting evidence across thousands of studies is impractical to perform manually. Recent advances in artificial intelligence (AI), particularly in large language models (LLMs), provide new opportunities to automate toxicological literature analysis.

**Results:** Herein, we present the Adverse Outcome Pathway Toolkit (AOPTK), an open-source Python package for AI-enhanced analysis of toxicological and biomedical literature. AOPTK supports (i) automated literature retrieval from databases such as PubMed Central and Europe PMC, (ii) direct processing of full-text articles in PDF format, and (iii) LLM-assisted analysis tools to identify chemicals and extract their relationships to biological effects under user-defined conditions. We demonstrate its utilities through a case-study identifying potential non-genotoxic carcinogens based on their ability to inhibit gap junction intercellular communication (GJIC). The LLM-enhanced analysis achieves 81% agreement with an expert-curated inhibitor list and 95% accuracy in predicting inhibition versus no inhibition when analysing the same subset of 86 publications. Automated retrieval and analysis of 1 124 publications from PubMed Central further demonstrated the feasibility of applying AOPTK to large-scale literature analyses.

**Availability:** The package is developed open source on GitHub, disseminated through PyPI and Bioconda and releases are archived on Zenodo. Individual tools are available in the Galaxy platform for accessible data analysis with user-friendly, findable, accessible, interoperable, and reusable (FAIR) and end-to-end workflows provided via WorkflowHub. The tools are available on usegalaxy.eu and usegalaxy.cz and are developed open source on GitHub.

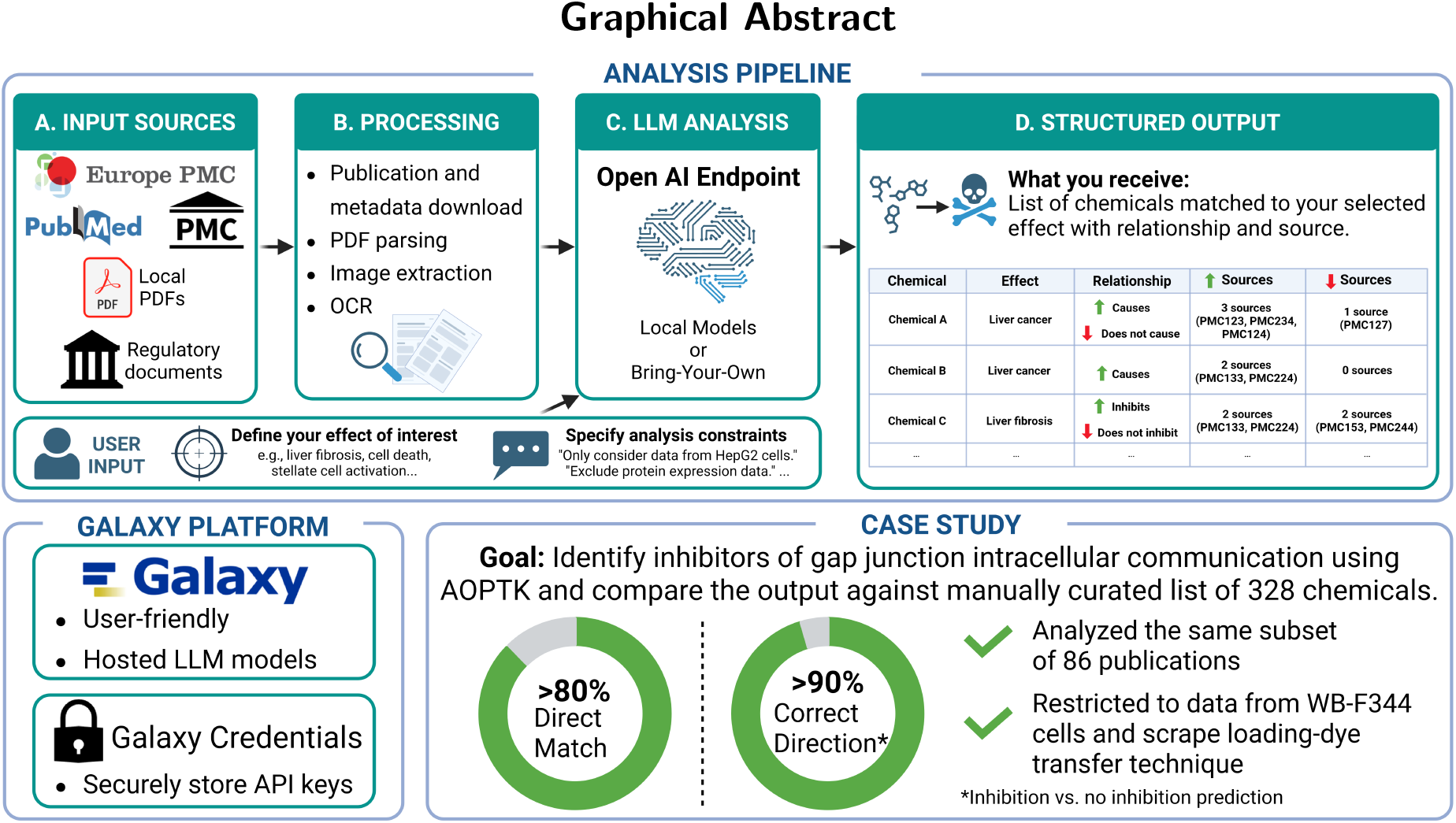

## Introduction

The number of scientific publications is persistently growing at an un-precedented rate (Bornmann et al., 2021), making systematic manual screening at scale increasingly impractical. In toxicology and bio-medicine, substantial amounts of valuable data remain hidden within unstructured text across publications, limiting their accessibility and reuse for scientific research, risk assessment of chemicals and regulatory decision-making (Hoffmann et al., 2017; Bajard et al., 2023). Recent advances in artificial intelligence (AI), particularly regarding large language models (LLMs), have demonstrated a remarkable ability to process, summarize, and extract structured information from vast corpora of unstructured scientific data, enabling new approaches to knowledge discovery in life sciences (Schmallenbach et al., 2024; Wang and Zhang, 2024).

In mechanistic toxicology, automated extraction of knowledge from textual data has already demonstrated considerable potential (Scholten et al., 2022; Bachman et al., 2023; Corradi et al., 2024), especially for chemical named entity recognition (NER) (Eltyeb and Salim, 2014; Neumann et al., 2019; Islamaj et al., 2021; Tarasova et al., 2022) and development and refinement of Adverse Outcome Pathways (AOPs) (Jaylet et al., 2025; Kumar et al., 2026), which provide structured representations of biological key events (KEs) linking molecular-level perturbations to adverse health outcomes. Furthermore, LLMs have emerged as a promising tool for knowledge extraction (Kumar et al., 2024; Luo et al., 2024; Djidrovski et al., 2026; Houweling et al., 2026; Na and Kim, 2026) as they offer improved contextual understanding, enhancing both accuracy and scalability of automated processes.

Herein, we present the AOP-toolkit (AOPTK). Originally developed to support the construction of AOPs, the toolkit provides generalpurpose functionality for the retrieval, processing, and LLM-driven analysis of scientific texts. Although a number of tools have successfully addressed individual aspects of literature retrieval and parsing (Swain and Cole, 2016; Bachman et al., 2023; Moreira-Filho et al., 2025), combining these capabilities into an interoperable and maintainable framework remains challenging. Scientific literature is distributed across multiple locations and file formats on the web, each requiring dedicated parsing approaches. The extraction of structured information such as tables, figures, and metadata is often complicated by inconsistent document formatting and limited machine readability, while database-specific application programming interface (API) restrictions can affect large-scale data collection. AOPTK addresses this need by integrating automated publication retrieval from multiple scientific databases, PDF parsing, chemical NER and AI-powered text analysis within a single framework following FAIR research software principles (Barker et al., 2022). In particular, this enables the automated extraction of chemical–biological effect relationships at scale (i.e., facilitating the linkage of stressors/chemicals to KEs and adverse outcomes in the AOP framework) while maintaining traceability to the underlying evidence. In addition, AOPTK supports the harmonization of chemical names across publications, ensuring consistency of the extracted data and matching of results across an entire corpus of texts. The toolkit is available on the Galaxy platform (The Galaxy Community et al., 2026), an open-source, web-based scientific workbench that enables complex computational data analyses through an intuitive graphical user interface (GUI), without requiring programming expertise or access to computational resources, while ensuring full reproducibility by automatically tracking all parameters, tools, and data provenance, making the toolkit’s capabilities (including high-end open-weight models) accessible to both technical and non-technical users.

## AOP-toolkit (AOPTK)

### Workflow

The AOPTK software package provides a comprehensive multistep pipeline for analysing scientific publications, including (i) data retrieval, (ii) chemical NER, (iii) chemical-effect relationship identification, (iv) chemical name normalization and (v) aggregation and alignment of results, as illustrated in Figures 1 and 3. Starting from a user-defined query term (e.g. “gap junction intercellular communication”) it supports the collection of abstracts and full text articles as plain text or as PDF from various literature databases, including Europe PMC, PubMed Central and PubMed. Alternatively, users can provide their own documents in PDF format for processing (Figure 1). The process of downloading publications consists of several steps. First, a selected database is queried using the user-defined search term to retrieve a list of relevant identifiers (IDs). During this step, Biopython (Cock et al., 2009) is used to access NCBI resources, such as PubMed Central, while Europe PMC is queried directly through its API. Second, the publications are downloaded by providing the ID list to any of the database clients. Note that due to technical limitations, not all combinations of databases and resources are supported. Each publication is then processed and analysed independently, ensuring that analyses remain self-contained. The analysis operates in two stages. First, chemical entities are identified; second, the text is analysed to extract relationships (e.g., causation, inhibition, …) between the recognized chemical entities and a user-defined biological effect. Those analyses can be tailored towards a specific domain or application through inclusion of user-defined free text (e.g., method specifications, restrictions to specific condition) in the prompts sent to the LLM.

**Figure 1.**
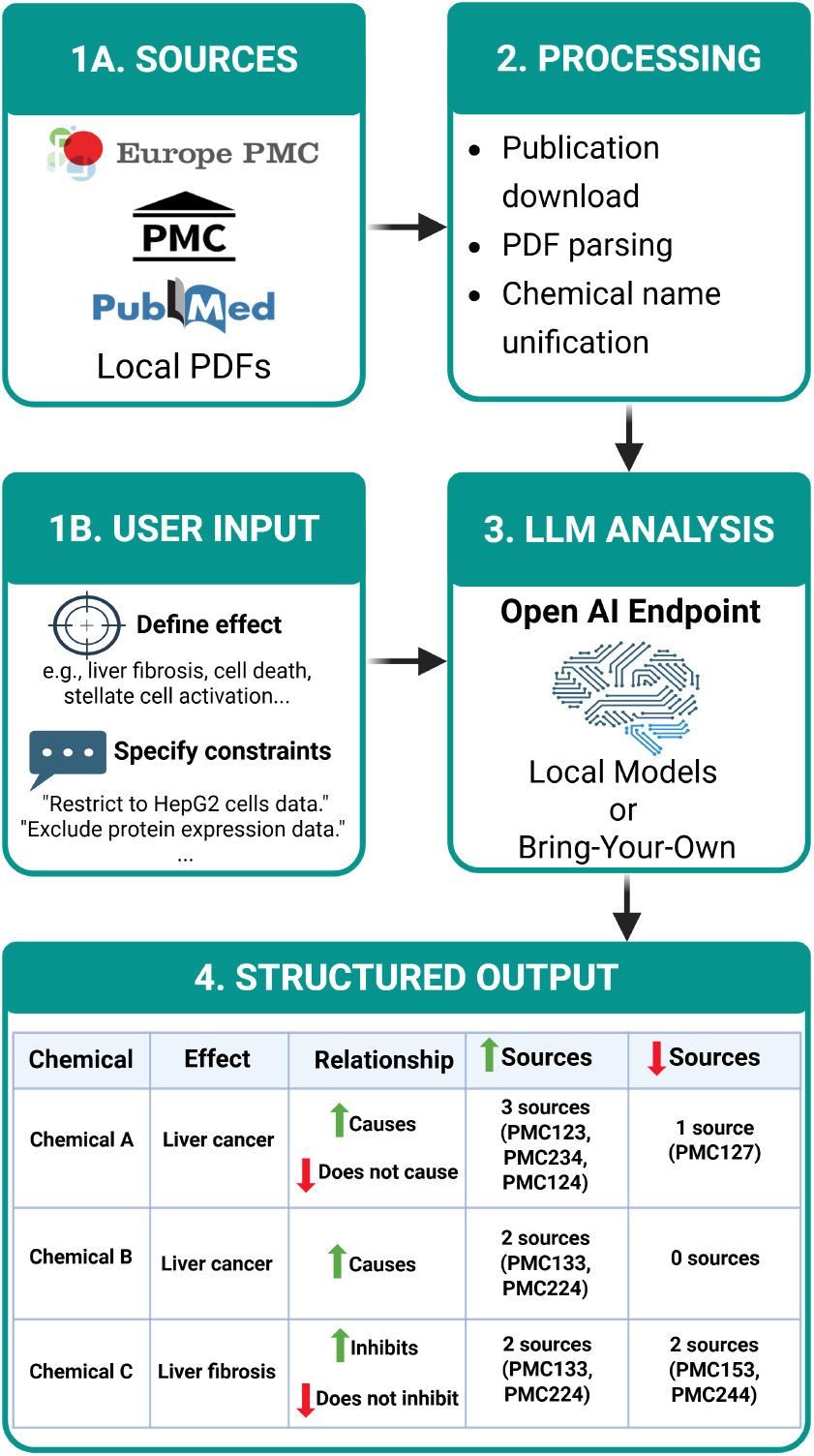
Overview of the AOPTK analysis workflow. Data is collected from various databases and locally stored PDF documents. Users define the effect of interest and can specify analysis constraints. The processed literature is then analysed using LLMs accessed via the OpenAI endpoint. A structured output containing extracted chemical–effect relationships is subsequently generated. For each chemical, the output reports the observed effect, the inferred relationship, and the number of literature sources supporting or contradicting the relationship, together with the associated reference IDs.

A major challenge in chemical-oriented literature analysis lies in the variability of terminology and naming conventions. It is common for a chemical substance to have various names used [e.g. consumer name (Aspirin), common name (Acetylsalicylic acid) and IUPAC name (2-acetyloxybenzoic acid)], making automatic matching across datasets increasingly difficult. Even though various structure-oriented aligned identifiers exist (i.e., InChI, SMILES), those are not commonly and systematically used throughout diverse scientific literature. Thus, after identification of chemicals and effect-relationships in individual publications, chemical names obtained through NER must be normalized prior to analysis of the whole corpus. AOPTK provides functions to normalize chemical names across publications via PubChem synonyms lookup, MeSH terms, or by matching terms against a user provided chemical name list using LLMs. It should be noted that the scope of this functionality goes beyond the capabilities of AOPTK and is a topic for future investigation but integration with the Galaxy platform allows direct post-processing with other relevant tools (Bray et al., 2020; Troják et al., 2022).

After normalization, the results from the individual analyses are combined to provide a summary of the chemicals associated with the biological effect, the type of relationship reported and the list of publications reporting the chemical-effect relationship. The ability of AOPTK to work with a broad range of effects – including KEs, diseases, molecular mechanisms, and other biological outcomes – supports diverse applications, such as identifying reference chemicals for an assay or KE/AOP, selecting appropriate assays for chemical testing, and identifying chemicals (including emerging chemicals) linked to KEs/AOPs.

### Implementation

The software is developed as an open-source project and is available on GitHub (Durník and Hecht, 2025). It is distributed via PyPI and Bioconda, ensuring easy installation. Comprehensive user, developer and API documentation is provided via Read the Docs. Furthermore, the GitHub repository includes a lightweight development container that provides a consistent developer environment, simplifying setup and streamlining contributor onboarding. Versioned releases are automatically archived on Zenodo (Durník and Hecht, 2026a) for improved findability and accessibility. To enhance overall software quality, SonarQube Cloud is employed for continuous AI code review and quality assessment using static code analysis. Further GitHub actions implement continuous testing to ensure functionality and maintainability of new source code additions (Figure 2). AI tools were used during software development. All generated artifacts underwent review by the authors before being integrated into the codebase. The AOPTK software is designed following object-oriented principles with a modular architecture that allows integration or replacement of different databases and analysis components, supporting maintainability and adaptability as models and data infrastructures evolve (Supplements 1, 2 and 3). Furthermore, this modular design opens possibilities for re-use in different domains. While the current implementation identifies relationships between chemicals and effects given a certain specification, the analysis framework revolves around rather abstract and simple statements in a subject (chemical) – predicate (relationship) – object (effect) – qualifier (specification) structure. The LLM-powered analysis engine is built on prompt templates which are populated with statements and context instantiations and leverages the OpenAI API endpoint specification, preparing the tool for variable deployments. Extending this framework towards other domains thus only requires implementation of equivalent statement classes at the same layer and the specification of prompt templates which provide the baseline instructions for the model.

**Figure 2.**
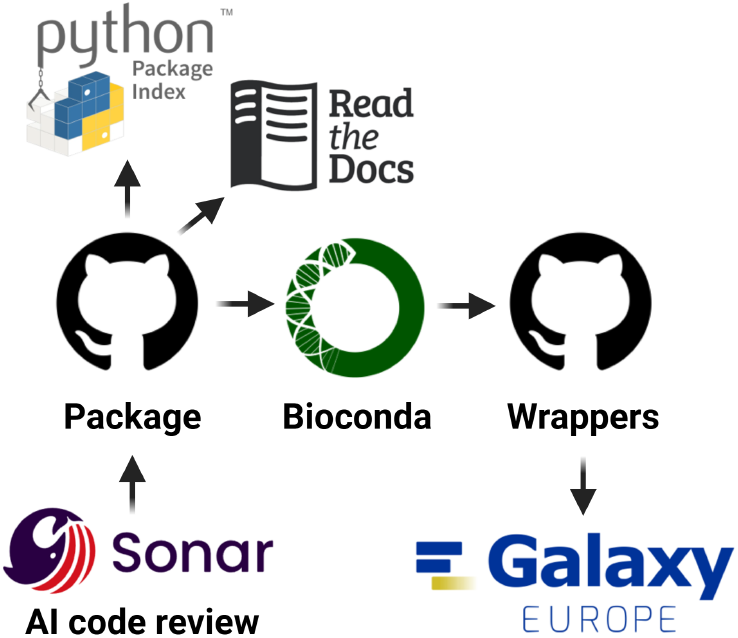
Technical stack behind the AOPTK software package that ensures the overall code quality, maintainability, documentation, and easy installation.

Galaxy tools wrapping the individual steps of the workflow (i.e. query literature, download articles, parse PDFs, analysis) are available on Galaxy Europe (The Galaxy Community et al., 2026; Hecht et al., 2026), providing a user-friendly web interface and making the tool accessible to a broader range of users (Figure 3). Via Galaxy, analyses can be exported as Research Object crates (RO-Crates) (Soiland-Reyes et al., 2022), which serve as reproducibility assurance: users can re-upload these crates back into Galaxy and fully reconstruct the analysis workflow, ensuring transparency and preventing fabrication. Besides providing a platform for accessible, reproducible and scalable analyses, open-weight LLMs deployed on research clouds such as eIn-fraCZ are accessible from multiple Galaxy servers (i.e., usegalaxy.eu and usegalaxy.cz). As of July 2026, the Galaxy platform supports various LLMs, including GPT OSS 120B and DeepSeek v3.2. Alternatively, users may integrate their own LLM services by securely storing the target OpenAI compliant API endpoint and authentication credentials in the Galaxy credentials system, enabling seamless access to external model deployments while securing secrets such as API keys.

**Figure 3.**
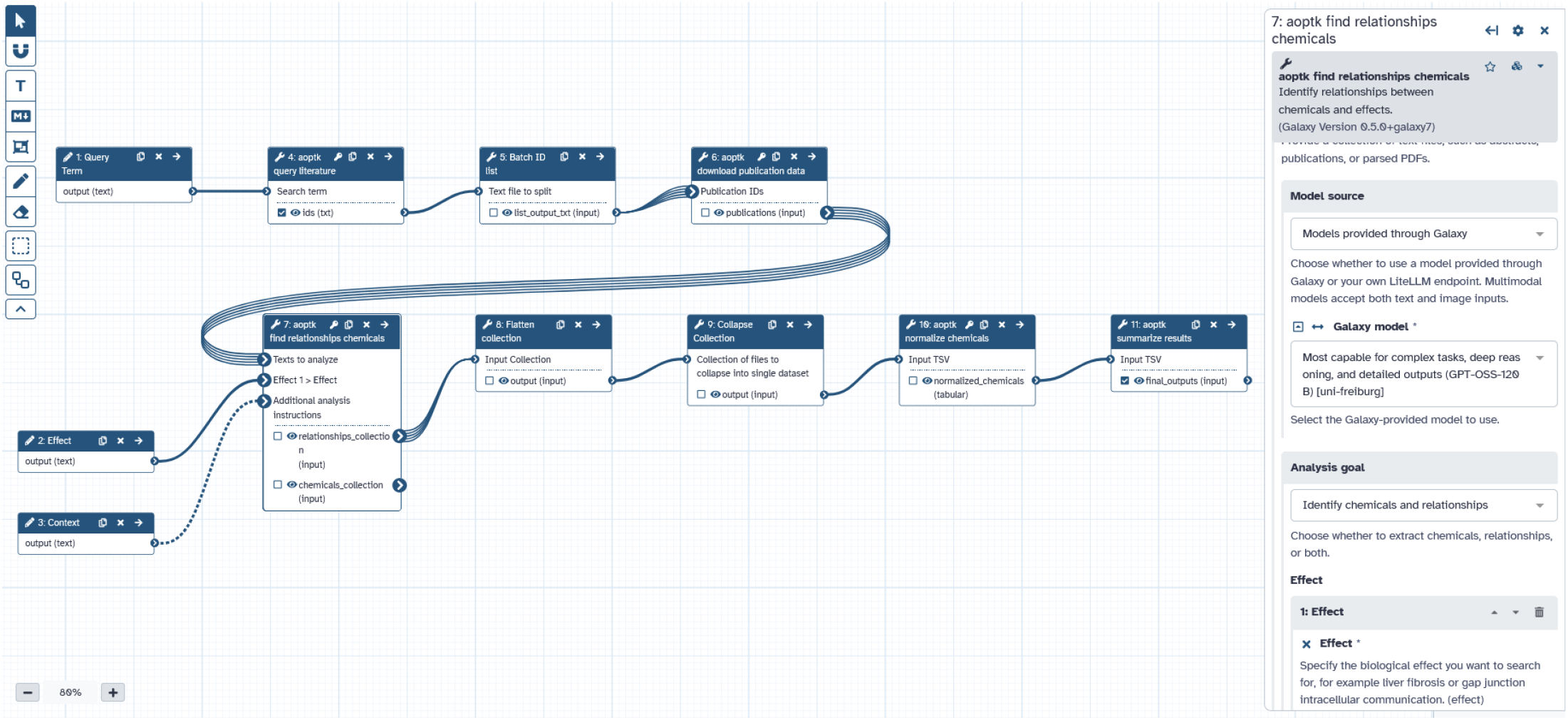
Galaxy workflow leveraging the AOPTK toolkit functions as various Galaxy tools. Publication IDs are first obtained and split into batches, which are then used to download the publication data. Publications are subsequently analysed using LLMs, and chemicals are harmonized across publications. The final list of chemicals is then provided to the user.

## Identification of chemicals inhibiting gap junction intercellular communication (GJIC)

To demonstrate its applicability, we present a case study using the AOPTK to identify chemicals associated with the inhibition of GJIC, a recognized hallmark of potential non-genotoxic carcinogens (Sovadinová et al., 2021).

### Case Study A: Comparison against expert-curated list of chemicals

To identify chemicals associated with inhibition of GJIC, we first benchmarked the toolkit output against a previously published list of 328 compounds manually curated by Sovadinová et al. (Sovadinová et al., 2021), which comprises both inhibiting and non-inhibiting substances. We used AOPTK to analyse the same set of 86 publications used in the original review (Supplements 4), provided in PDF format. In alignment with the review’s methodology, we restricted the analysis to data generated in WB-F344 cells using the scrape-loading fluorescent dye transfer assay. Relevant data was extracted from the publications using LLM-based prompting. We leveraged the GPT OSS 120B LLM to normalize the chemical names across publications and to match them to the expert-curated list.

The LLM-based analysis retrieved 81% of the chemicals from the list (266 chemicals matched out of 328 chemicals), and the direction of the effect (inhibition vs. no inhibition of GJIC) was correctly assigned in 95% of these cases (252 chemicals out of 266 assigned correctly). Results of the analysis, run on the Galaxy platform, were exported as RO-Crate and are available on Zenodo (Hecht and Durník, 2026). Exact comparison to the expert curated list can be found in Supplements 5. 158 extra chemicals (inhibiting and/or non-inhibiting), not present in the expert-curated list, were identified by the LLM. Numerical matches with the expert-curated list are represented by a Venn diagram (Figure 4).

**Figure 4.**
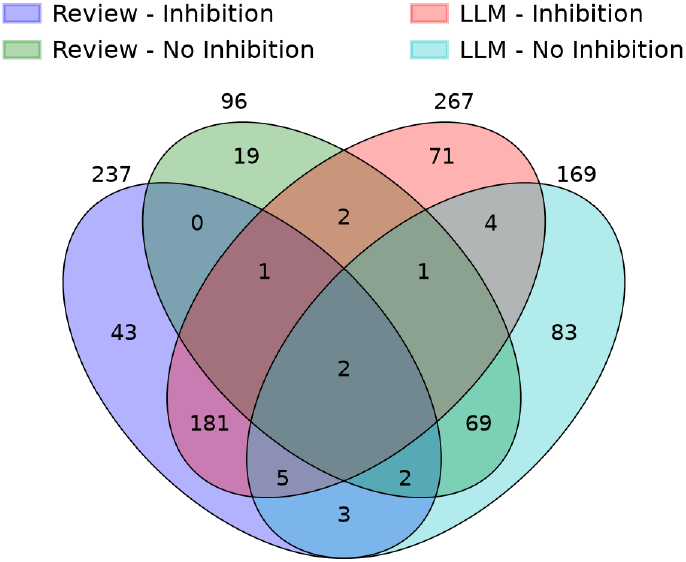
Venn diagram illustrates the overlap between manually curated (“Review”) and AOPTK-retrieved (“LLM”) lists of GJIC inhibitors when analyzing the same subset of 86 publications. For clarity of the graphical presentation, chemicals with mixed effects (both inhibition and no inhibition) were counted in both categories.

### Case Study B: Large-scale analysis of open access literature

As a second case study, we again applied the AOPTK to generate a list of chemicals associated with GJIC, this time screening the PubMed Central database. We systematically analysed the full texts of open-access publications retrieved using the following search term (1 124 records in PubMed Central):

gap junction intercellular communication inhibition [Title/Abstract] AND 2020/01/01:2026/07/08[dp]

Analysis was run again using the GPT OSS 120B model. The resulting list of chemicals associated with the inhibition of GJIC, harmonised using PubChem synonyms, is available in Supplements 6 and exported as RO-Crate. The analysis identified 3335 unique chemical structures described via InChI out of which 157 could be directly linked to inhibition (or no inhibition) of GJIC.

## Discussion

A key factor in software adoption is accessibility, particularly for users outside the technical community. This especially applies in the case presented here, as the primary target audience consists of toxicologists, many of whom may not have extensive technical expertise in informatics. AOPTK is a user-friendly and accessible software package that leverages LLMs to identify and extract chemicalbiological effect relationships from published studies, transforming unstructured text into structured knowledge that can support downstream analysis and decision-making. To maximize accessibility, we implemented AOPTK on the Galaxy platform, which provides a userfriendly web interface and integrates seamlessly with existing analysis workflows. Users can either leverage locally hosted LLMs available within Galaxy or connect their own models by securely storing API keys and endpoints through Galaxy credentials. We demonstrated the capabilities of AOPTK by generating lists of chemicals associated with GJIC inhibition.

### Case Studies

In the first scenario (case study A), we analysed a smaller subset of 86 publications available in PDF format, benchmarking the automated AOPTK approach against expert-derived results (Sovadinová et al., 2021). The performance for case study A (Supplements 5) achieved 81% direct match and 95% correct direction prediction (inhibition vs. no inhibition). The expert-derived list provided aggregated evaluations for each chemical, integrating evidence across multiple publications. Some chemicals were classified as having equivocal or mixed evidence rather than being assigned a simple binary outcome (e.g., anthracene and benzo[e]pyrene). Consequently, complete direct comparison was not always possible. The LLM workflow also identified a substantial number of chemicals that were absent from the expert list. Several factors explain these discrepancies. Some chemicals may not have been included in the original list, for example because they were used as positive or negative controls rather than being the primary focus of evaluation (e.g., acetonitrile). Additional chemicals may have been retrieved due to limitations in the prompting strategy, which may not have sufficiently constrained the extraction to the desired evidence. In practice, the tool is expected to be applied primarily to larger-scale case studies, where chemicals with only limited literature support would typically be filtered out using predefined thresholds. Chemicals missed by the LLM may have been reported only in figures, image-based tables, or other graphical content that was not analysed by the AOPTK workflow.

Nevertheless, concordance between manual curation and LLM-assisted extraction was generally high. For example, both approaches consistently identified several environmentally relevant chemicals associated with non-genotoxic modes of action, including DDT, lindane, pentachlorophenol, PFOS, and PFOA, as inhibitors of GJIC. Conversely, both approaches consistently classified compounds such as PFBS, PFPeA, octanoic acid, decanoic acid, propyl gallate, and epicatechin as non-inhibitory. These examples illustrate that the workflow can recover both positive and negative evidence relevant to GJIC disruption and may therefore support the identification of potential non-genotoxic carcinogens while also providing candidate true-negative reference compounds for assay validation. Chemicals with conflicting and inconclusive literature evidence, such as anthracene, benzo[e]pyrene, and dibenz[a,c]anthracene, were often represented by both inhibitory and non-inhibitory observations in the extracted dataset. The primary advantage of utilizing LLMs lies in their ability to scale beyond human reading capacity. The second case study thus primarily demonstrated the scalability of the workflow rather than providing a comprehensive assessment of GJIC-associated chemicals. In the case study B, we applied AOPTK to a larger corpus of open-access literature published from 2020 onward, to generate a new list of 157 chemicals associated with GJIC inhibition (Supplements 6) by retrieving and analyzing articles directly from PubMed Central. The recent literature is heavily dominated by a small set of well-established GJIC inhibitors. Carbenoxolone was by far the most frequently reported inhibitor (51 studies), followed by mefloquine, glycyrrhetinic acid derivatives (18*α*- and 18*β*glycyrrhetinic acid), meclofenamate/meclofenamic acid, quinine, tonabersat, Gap27/Gap19 peptides, and long-chain alcohols such as 1-octanol and heptanol. These compounds represent the most consistently validated pharmacological tools for inhibiting gap junction intercellular communication. Beyond these reference inhibitors, the dataset also includes several environmentally relevant chemicals with known or suspected carcinogenic properties, including bisphenol A, bisphenol S, hexachlorocyclohexane, DDT-related compounds, and short-chain chlorinated paraffins, which have been reported to inhibit GJIC in individual studies. These observations support further investigation of GJIC disruption as a mechanistically relevant endpoint for the identification of potential non-genotoxic carcinogens. Several compounds yielded more ambiguous results. Arachidonic acid and other polyunsaturated fatty acids, calcium signaling modulators, nitric oxide, glucose-related treatments, and oxidative stress-associated agents were reported as inhibitors only in some studies, suggesting that their effects may depend on cell type, concentration, exposure conditions, and underlying signalling pathways. Overall, the evidence indicates a clear distinction between a relatively small group of robust and widely accepted GJIC inhibitors, which may serve as positive reference compounds for assay development, optimization, and inter-laboratory validation, and a much larger group of compounds whose effects on gap junction communication remain conditional, indirect, or insufficiently validated. In practice, such large-scale analyses could support two important applications:

- Identification of well-supported chemicals (i.e., those with a high number of literature mentions), which may be useful for method validation or updating existing chemical lists (e.g., carbenoxolone in our case study).
- Pre-filtering of emerging or under-reported chemicals (chemicals with low number of literature mentions), which can serve as candidates for further manual text mining or experimental (toxicological testing) validation. This is an important step to prioritize chemicals for hazard and risk assessment.

Working with chemical data introduces several inherent limitations – a major one being the variability in chemical nomenclature. To mitigate this, AOPTK employs multiple approaches to harmonize chemical names. This includes PubChem synonym lookup or LLMs (matching a chemical name against a user-provided chemical list). This limitation affects both the final tool output – where identical chemicals may be represented under different names – and comparisons with expert-curated lists. Consequently, the observed match ratios in our analysis may be underestimated. Beyond the nomenclature, the NER component is not strictly limited to chemical structures, but also recognizes groups of chemicals (e.g. alcohols, pesticides, PCBs) which comprise multiple structures or other mixtures which are grouped under a single common name. Matching those entities against another list poses several problems as equivalence might be variable or not well-defined. Additionally, structural isomers present a particular challenge, as LLMs often struggle to identify and distinguish between them. This issue is further complicated by scientific literature itself, where isomers are not always clearly distinguished. As a result, ambiguities in the source publications can be carried over into the software’s output. These limitations could potentially be mitigated through improved prompting strategies or by leveraging more advanced LLMs as they become available.

### Implementation

The ability to analyse locally stored PDF documents offers several important advantages. Firstly, it reduces dependence on scientific literature databases by enabling the direct processing of publications. Secondly, it broadens the scope of analysis beyond scientific articles to potentially include any other text resources such as regulatory documents. Despite these advantages, several limitations remain. Parsing publications in PDF format is inherently challenging due to inconsistent formatting and accessibility issues. Older publications are often available only as scanned documents rather than embedded text PDFs. In such cases, multimodal LLM capabilities can be used to extract and convert the content into machine-readable text for subsequent analysis. Tables present an additional difficulty, as they may appear either as embedded text or as images, leading to inconsistent extraction performance. While image extraction itself is relatively reliable, downstream automated analysis of visual content remains limited due to current constraints of LLM capabilities and thus was not performed in this study.

The analysis is conducted initially at the level of individual publications and then aggregated by matching across the entire literature corpus. This approach preserves the full granularity of the underlying evidence and avoids oversimplification, as the tool goes beyond merely summarizing findings across multiple publications. Users can easily identify whether both positive and negative relationships have been assigned to the same chemical and can therefore decide to classify the results as equivocal. Furthermore, because the analysis processes entire publications, results extracted from introduction and discussion sections – such as references to findings from other studies – are also included, rather than only novel experimental outcomes. This can lead to an over-representation of certain chemicals. One improvement would be to restrict analysis to experimental result sections, either through refined prompting or improved parsing of publication structure. However, this presents a technical challenge, as publications vary substantially in format, making consistent extraction of specific sections across different articles difficult. Due to the stochastic nature of LLMs, outputs may vary between analysis runs even when the same input data and prompts are used.

## Conclusion

AOPTK addresses a key need of the toxicology community by substantially reducing the time and effort required to generate literature-based reviews and chemical lists. As a FAIR, open-source research software framework, it enables transparent, reproducible, and scalable extraction of toxicological knowledge from scientific literature. AOPTK is intended to complement the expert assessment by providing a quick, evidence-based starting point that guides experts toward relevant chemicals, mechanisms, and publications for further evaluation. By enabling systematic reuse of existing evidence, it can accelerate chemical safety assessment and support a wide range of applications in toxicological research. As demonstrated in this research, AI can provide toxicologists with powerful tools to deal with and efficiently re-use the data they have already generated.

## Data Availability Statement

The data underlying this article is available on Zenodo (Hecht and Durník, 2026). The main software is developed open source on GitHub at https://github.com/RECETOX/aoptk and releases are archived on Zenodo for preservation (Durník and Hecht, 2026a). The galaxy tools are also developed open source and are available on GitHub at https://github.com/RECETOX/galaxytools and are archived on Zenodo (Hecht et al., 2026). The presented workflow is freely available on usegalaxy.eu at https://usegalaxy.eu/u/hechth/w/aoptk-download-and-analysis and stable versions archived on WorkflowHub (Durník and Hecht, 2026b).

## Authorship

Robin Durník developed the software, performed the formal analysis, wrote the initial draft and contributed to the conceptualization. Lola Bajard contributed to the conceptualization, and formal analysis and contributed to review and editing of the manuscript. Luděk Bláha lead the conceptualization and contributed to review and editing of the manuscript. Pavel Babica supervised the formal analysis and contributed to review and editing of the manuscript. Iva Sovadinová supervised the formal analysis and contributed to review and editing of the manuscript. Helge Hecht lead the methodology, supervised the development of the software and contributed to the formal analysis, conceptualization and review and editing of the final manuscript.

