## Supplementary figures and images for "AOPTK: Open-Source Software for AI-Assisted Toxicological Literature Analysis"

### Figure S1

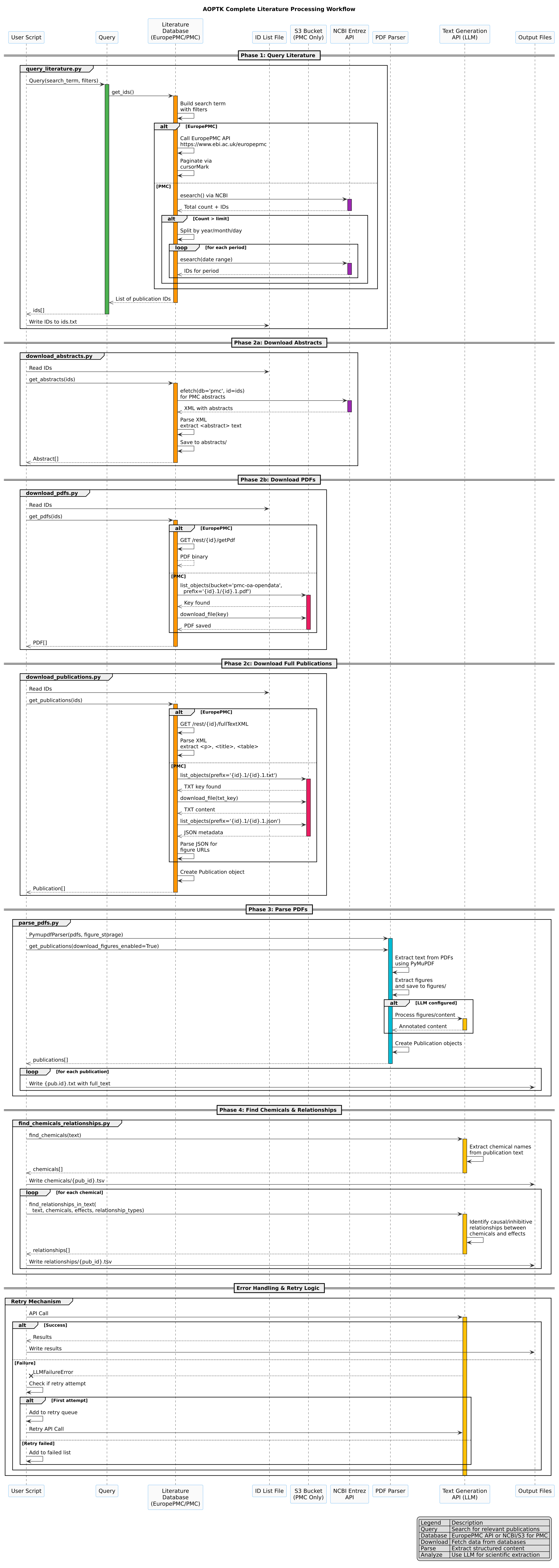

### Figure S2

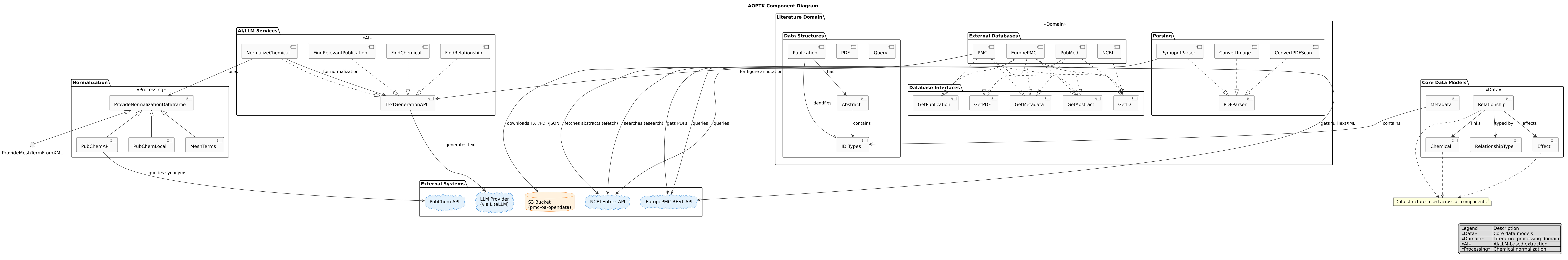

### Figure S3

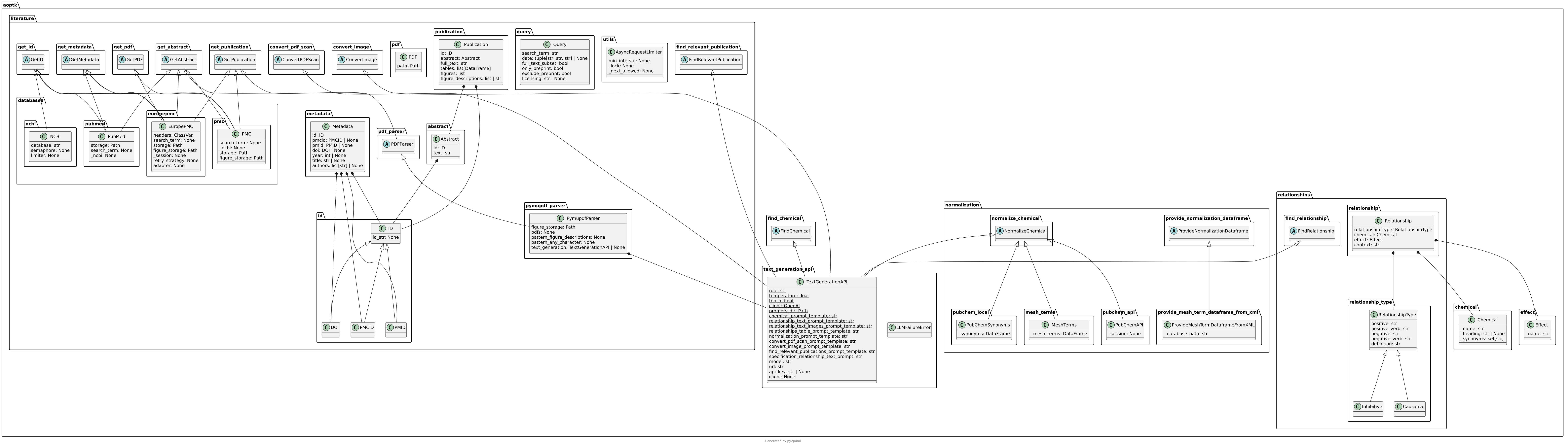
